# Nanoparticles co-delivering siRNA and mRNA for simultaneous restoration and silencing of gene/protein expression *in vitro* and *in vivo*

**DOI:** 10.1101/2024.06.22.600196

**Authors:** Shireesha Manturthi, Sara El-Sahli, Yuxia Bo, Emma Durocher, Melanie Kirkby, Alyanna Popatia, Karan Mediratta, Redaet Daniel, Seung-Hwan Lee, Umar Iqbal, Marceline Côté, Lisheng Wang, Suresh Gadde

## Abstract

RNA-based agents such as siRNA, miRNA, and mRNA can selectively manipulate gene expression/proteins and have the potential to revolutionize the current therapeutic strategies for various diseases, including cancer. To address the poor stability and inherent limitations of RNA agents, nanoparticle (NP) platforms have been developed to deliver functional mRNA or siRNA inside the cells. Recent studies have focused on either siRNA to knock down proteins causing drug resistance or mRNA technology to introduce tumor suppressors. However, complex diseases like cancer need multi-targeted approaches to selectively target multiple gene expressions/proteins. In this proof-of-concept study, we developed co-delivery nanoparticles containing Luc-mRNA and siRNA-GFP as model RNA agents ((M+S)-NPs) and assessed their effects *in vitro* and *in vivo*. Our studies show that NPs can effectively deliver both functional mRNA and siRNA together, simultaneously impacting the expression of two genes/proteins *in vitro*. Additionally, after *in vivo* administration, co-delivery NPs successfully knocked down GFP while introducing luciferase in a TNBC mouse model, indicating our NPs have the potential to develop RNA-based anticancer therapeutics. These studies pave the way to develop RNA-based, multitargeted, multi-delivery approaches for complex diseases like cancer.

**TOC:** 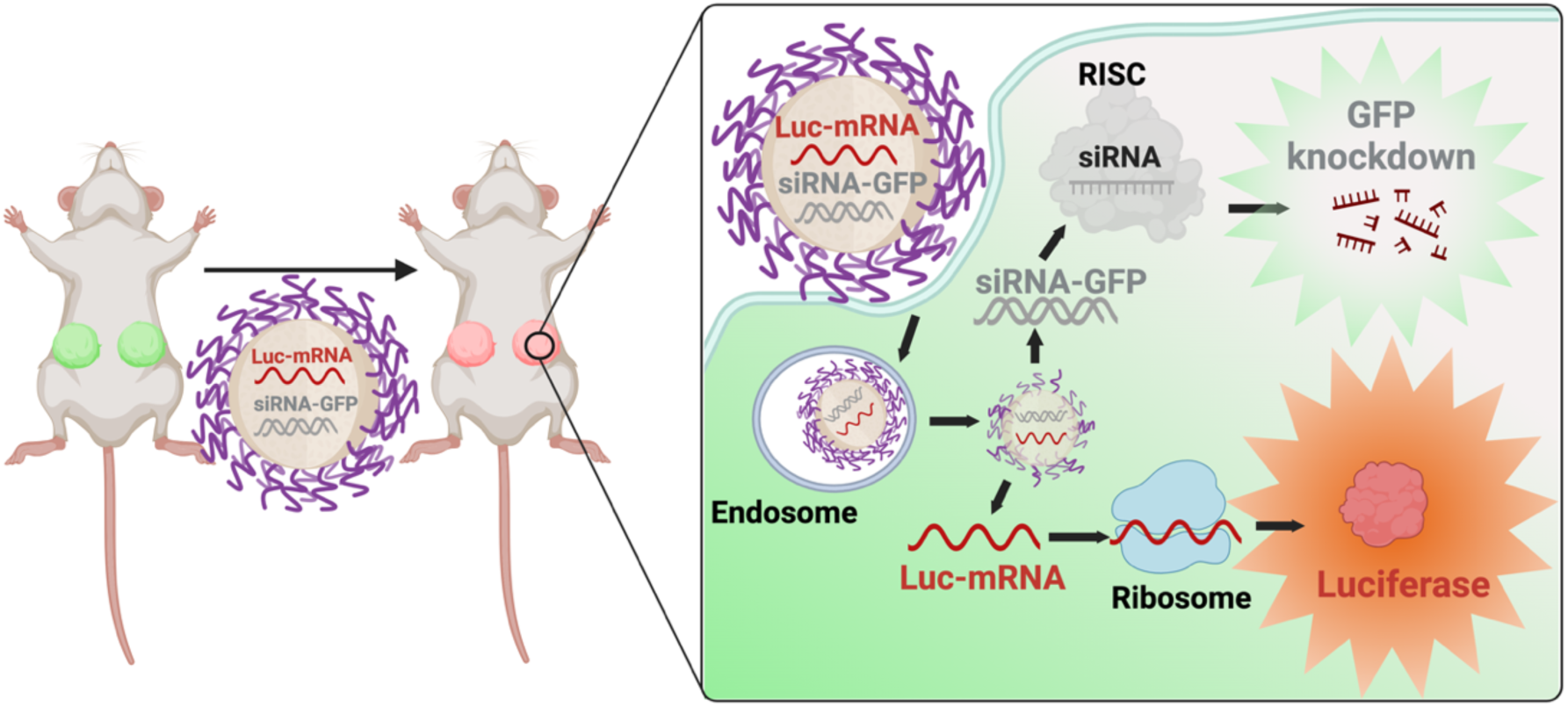

## Introduction

RNA-based therapies have the capacity to selectively manipulate gene expressions and hold the potential to revolutionize the current therapeutic strategies for various diseases, including cancer.^1–3^ RNA-based agents such as siRNA, miRNA, and mRNA can downregulate, augment, or correct specific gene products that are otherwise undruggable with small molecules.^4, 5^ mRNA agents, typically over 2000 nucleotides long, include conventional mRNA, self-amplifying mRNA (saRNA), trans-amplifying mRNA (taRNA), and circular mRNA (circRNA).^6^ These mRNA agents can be designed to carry optimized genetic information and be translated into the production of encoded proteins without integrating into the host genome.^1, 6–8^ mRNA technology offers greater flexibility for targeting various diseases, including cancer, cardiovascular diseases, as well as vaccines.^6, 8, 9^ In cancer treatment, mRNA applications are multitude. They can encode tumor suppressors to inhibit cancer cell proliferation, tumor antigens to trigger immune response, and chimeric antigen receptors (CARs) or T cell receptors (TCRs) for T cell therapies.^6, 10^ While mRNA therapies can produce specific proteins in the target cells, RNA interference (RNAi), a natural defense mechanism against exogenous genes, can specifically knock down target gene expression.^11, 12^ RNAi agents, such as siRNA and miRNA, modulate target genes by mediating targeted mRNA degradation (siRNA and miRNA) or mRNA translation repression (miRNA). Both siRNA and miRNA-based RNAi therapies have shown significant potential in cancer therapies, specifically in knocking down/modulating the expression of genes/proteins involved in drug resistance and cancer stem cell (CSC) enrichments, inducing cell death, and sensitizing cancer cells to chemotherapeutic drugs.^13, 14^

Despite their potential, mRNA and RNAi therapies face several challenges in clinical applications.^3^ RNA agents, such as mRNA, siRNA, and miRNA, possess undesirable physicochemical and pharmacological properties, are susceptible to degradation and unwanted immune reactions, and must be effectively delivered into the target cells.^2, 3, 14^ Therefore, sophisticated delivery platforms are essential for developing safe and effective nucleic acid-based therapies.^15^ Nanotherapeutic strategies offer several advantages over conventional therapies. Nanoparticle-based drug delivery platforms can encapsulate various therapeutic agents irrespective of their physiochemical properties, protect them from degradations, and deliver them into the cells.^16^ However, delivering RNA agents is more challenging than traditional therapeutic agents due to the large size of mRNA (over 2000 nucleotides) and the phosphate backbone common to all RNA agents, which complicates their encapsulation in NPs.^7, 11, 15, 16^ Additionally, NPs must escape the endosome and release functional RNA agents into the cytosol to be effective. To overcome these challenges, advanced NPs platforms will be required.^2, 11^ In this context, recent lipid NPs have shown success in delivering either mRNA or siRNA, with examples including Onpattro (patisiran), an siRNA-containing NPs currently in the clinic for treating hereditary amyloidogenic transthyretin (TTR) amyloidosis, and the Moderna Spikevax and Pfizer-BioNTech COVID-19 vaccines, which are NPs containing mRNA encoding the SARS-CoV-2 spike glycoprotein.^8, 17^ Despite these successes, strategies targeting a single protein or cytokine are insufficient for complex diseases like cancer and cardiovascular diseases, the leading causes of mortality worldwide.^18^

In cancer treatment, nanotherapies have been reported to knock down proteins involved in drug resistance or restore tumor suppressor genes to control tumor growth.^19, 20^ However, in cases like triple-negative breast cancer (TNBC), which accounts for a disproportionate number of breast cancer-related deaths, treatment strategies need to address significant challenges such as drug resistance, cancer stem cell enrichment, and tumorigenesis, representing an unmet medical need.^21^

We rationalized that combining mRNA and RNAi technologies could leverage their respective advantages to develop potent multi-targeted therapies for cancer and other diseases. Specifically, we envisioned that selectively restoring the expression of anti-cancer genes while knocking down pro-cancer genes would be an effective therapeutic strategies strategy. To achieve this, we developed a co-delivery NP system containing both mRNA and siRNA to simultaneously knock down and restore two different gene expressions. As a proof-of-concept, we used Luc-mRNA and siRNA against GFP, studying their effects *in vitro* and *in vivo* using GFP+ cells as a model. Our results demonstrated that NPs successfully encapsulated both mRNA and siRNA, protected them from degradation, delivered functional agents into the cells, and simultaneously and effectively knocked down and restored the target gene expressions in both *in vitro* and *in vivo* settings.

## Results

### Development and characterization of co-delivery NPs

For this proof-of-concept study, we first developed single and dual RNA agents containing NPs *via* a self-assembling process using PLGA_10K_-PEG_5K_ polymer. PLGA-PEG polymeric NPs were chosen due to their biodegradability, biocompatibility, stability, scalability, and versatility in various drug delivery applications.^22^ To address the hydrophilic nature of RNA’s phosphate backbone and facilitate encapsulation within the NPs, we used PEI-C_14_ as a cationic lipid (Figure S1&2). Both single and dual-drug NPs were synthesized using the nanoprecipitation method by blending siRNA and/or mRNA with PEI-C_14_, followed by the addition of PLGA_10K_-PEG_5K_, and dropwise addition to nuclease-free water.^21^ The resulting NPs were Luc-mRNA containing (M)-NPs, siRNA-GFP containing (S)-NPs, and Luc-mRNA + siRNA-GFP containing (M+S)-NPs (Figure 1). NPs were instantly formed with PEI-C_14_:siRNA and PEI-C_14_:mRNA complexes embedded in the PLGA hydrophobic core stabilized by a PEG shell.^23^

**Figure 1:**
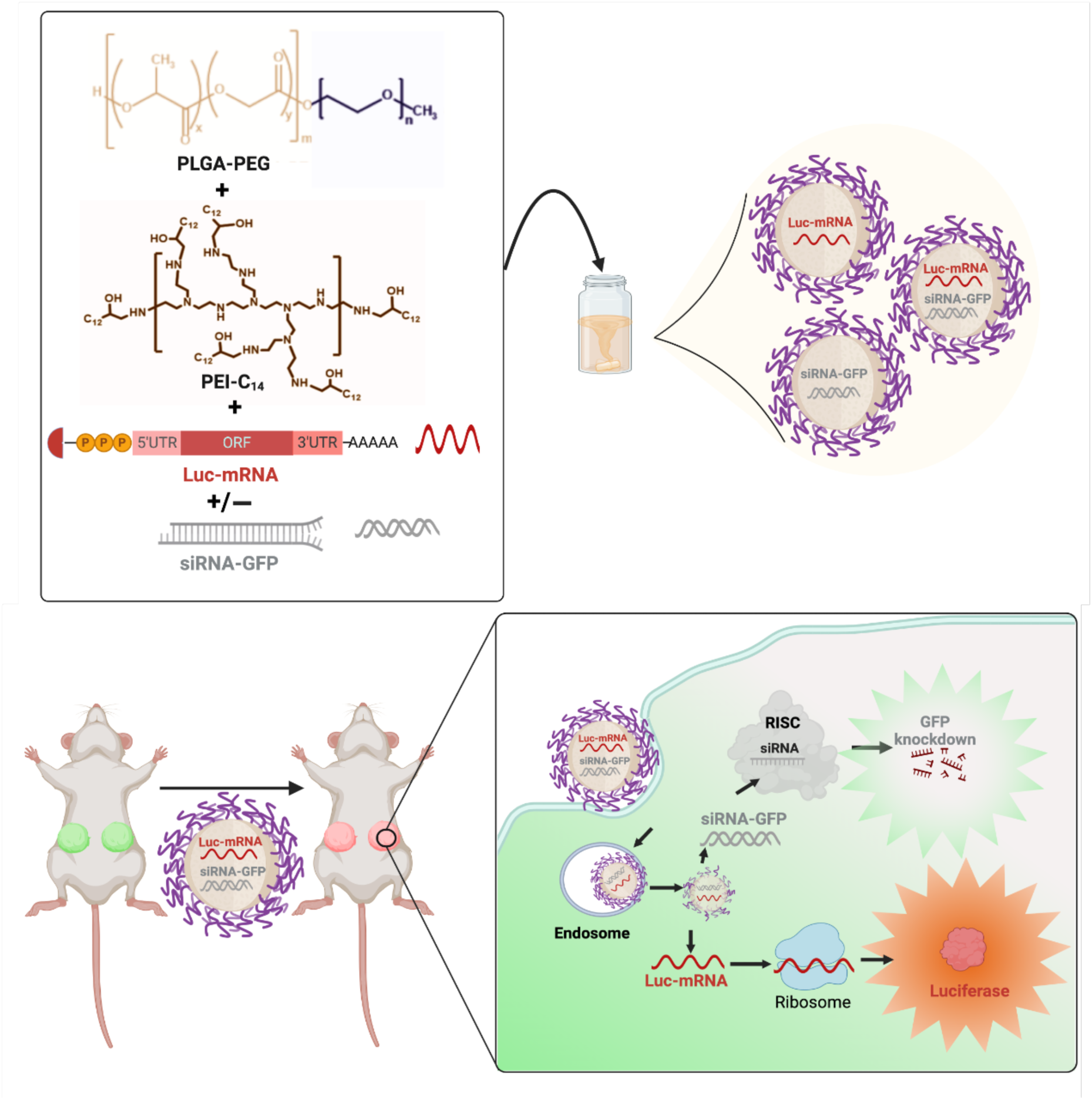
Schematic diagram of NPs development (Upper panel) and their *in vitro* and *in vivo* evaluation (Lower panel)

After synthesis, NPs were collected and purified using centrifugal filters and characterized for their physicochemical properties. Hydrodynamic size and surface charge were measured using dynamic light scattering (DLS). All 3 NPs have uniform sizes and range between 50 and 60 nm, with low PDI (0.18-0.25) and surface charges between −0.2 and −20 mV (Figure 2A-B, and E). TEM characterization of NPs showed all 3 NP formulations have a spherical shape, and the actual sizes are 40-50 nm, which are lower than DLS sizes as expected (Figure 2C). The % of encapsulation efficiency (EE%) of siRNA, measured using Cy5 labeled siRNA, was approximately 92% and 85% for single and dual drug NPs, respectively. EE of Luc-mRNA, determined by RiboGreen assay, was 100% and 91% for single and dual drug NPs, respectively.^24^ We then studied the serum stability of our NPs by incubating them 6 hours at different serum concentrations (up to 10%). Results indicated that NPs remained stable in the presence of serum proteins, as there were no significant changes in NP sizes before and after incubation (Figure 2D).^25^

**Figure 2:**
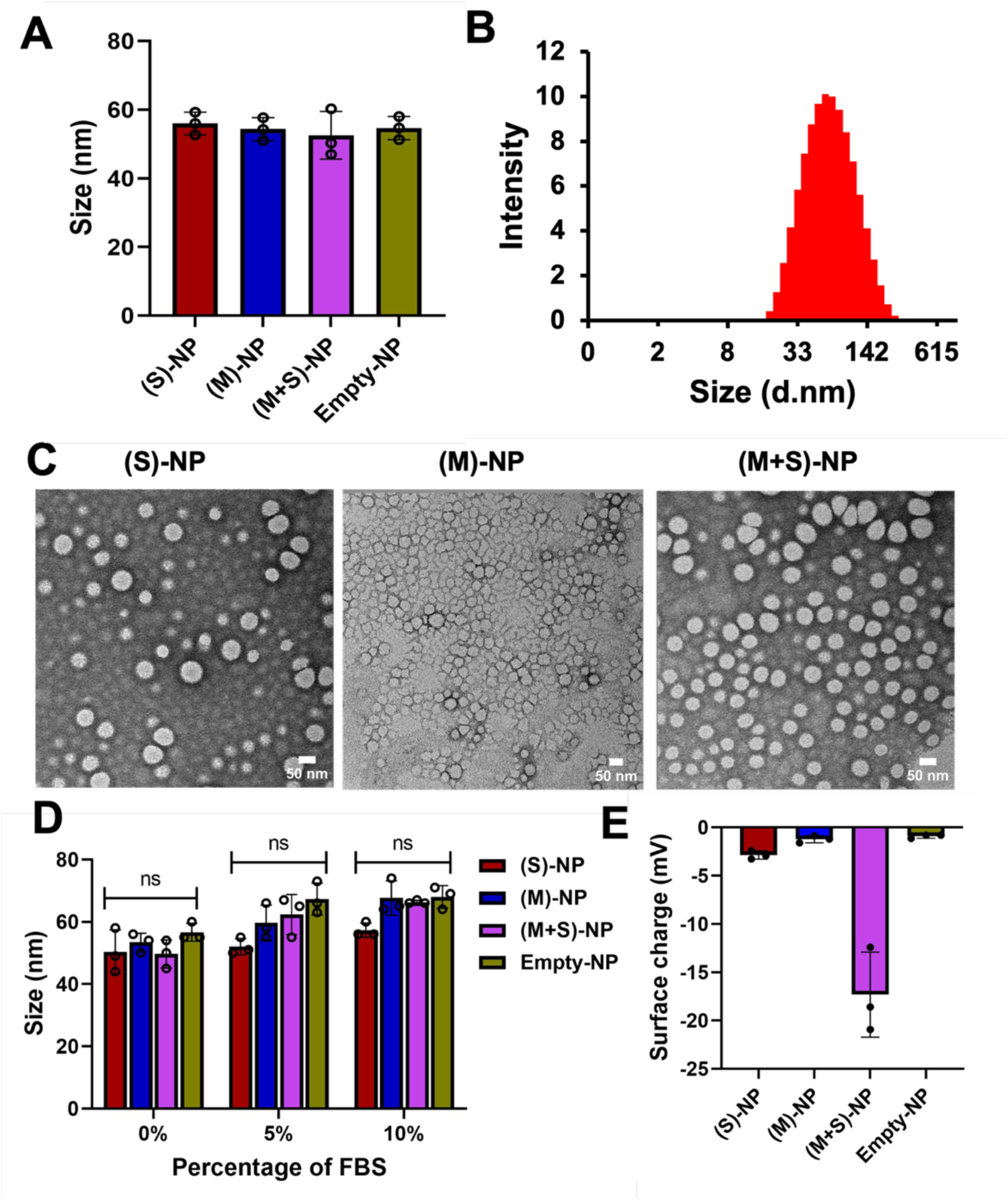
Physicochemical characterization of nanoparticles. (A) Size of single and dual-drug NPs along with control-NPs, measured by diluting 20 µL of NP in 1 mL sterile water using Dynamic light scattering n=3. (B) Size distribution intensity of (M+S)-NPs, measured by DLS. (C) Size and morphology of Single and dual-drug NPs by transmission electron microscopy (scale bar-50 nm). (D) Stability of all the NPs in different percentage of FBS; NPs incubated for 6 hours and measured the size; n=3 (ns-no significant difference). (E) Surface of single and dual-drug NPs along with control-NPs, measured by diluting 20 µL of NP in 1 mL using Dynamic light scattering n=3.

### Co-delivery NPs capable of protecting and delivering both functional mRNA and siRNA in vitro

To evaluate our NP’s capacity to escape endosomes and deliver functional mRNA and/or siRNA, we performed a flow cytometry study. For this purpose, we developed NPs loaded with Cy5-labeled siRNA to mimic siRNA-GFP and EGFP mRNA as model mRNA (Cy5+EGFP mRNA)-NPs). The co-delivery NPs size and surface charges are similar to (M+S)-NPs (S3 A-D). HT1080 cells were treated with (Cy5+EGFP mRNA)-NPs and control NPs for 48 h, and NPs delivery capacity was analyzed by flow cytometry. The Cy5 label could track siRNA in NPs, while GFP expression from the cells revealed functional mRNA delivery.^19, 26^

After 48 h treatment, we observed a significant percentage of cells associated with Cy5 (100%) (Figure 3B &C). and GFP (25%) (Figure 3D, E, &F) signals, demonstrating that NPs can deliver agents intracellularly and they are functional. Despite some Cy5 signal bleeding into the GFP channel, there was a 3-folds increase in GFP signal in the cells. These results indicate that the NPs successfully entered the cells and delivered both mRNA and siRNA.

**Figure 3:**
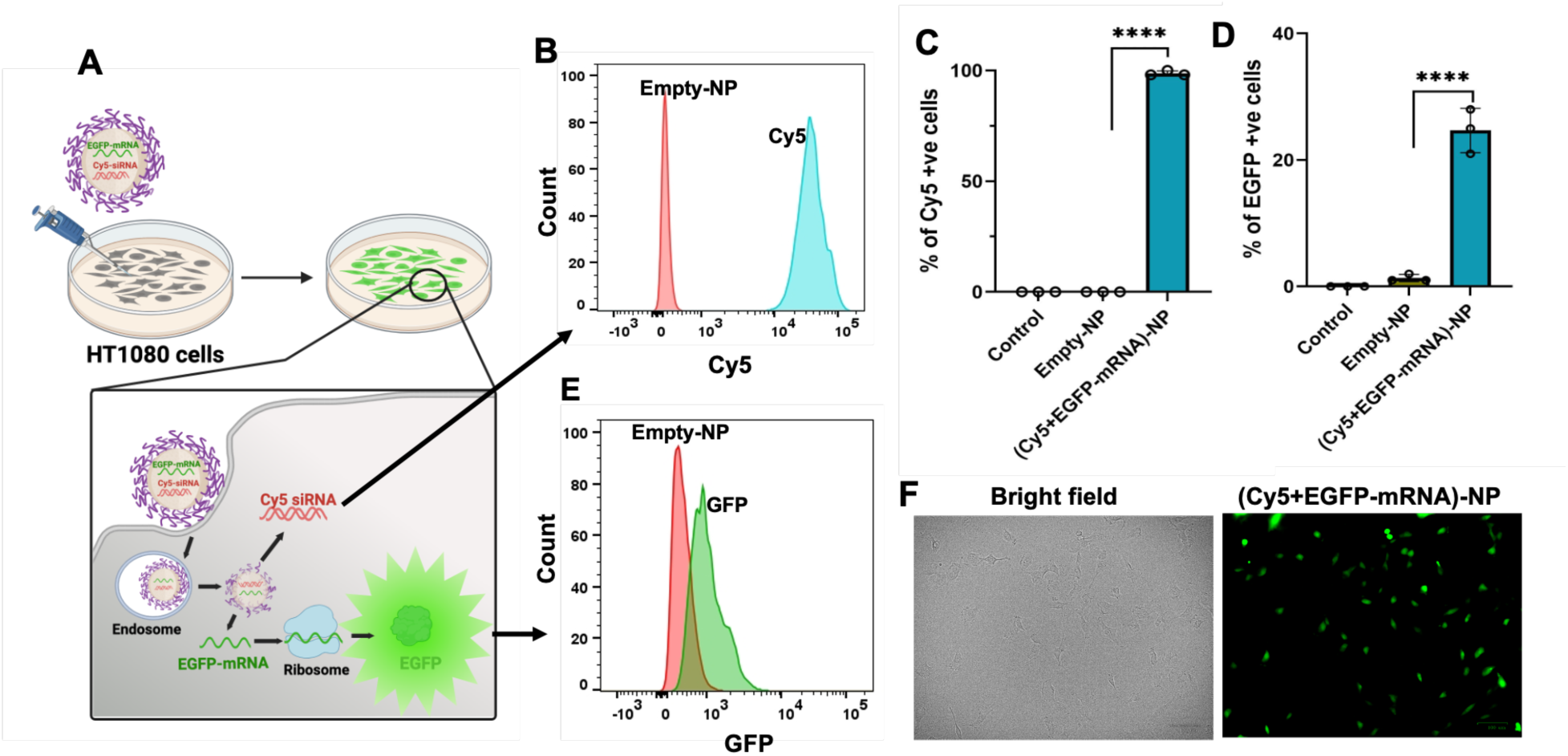
Cellular interactions of co-delivery NPs. (A) Model representation of NPs treatment to HT1080 cells; Cy5 cellular entry and expression of EGFP. (B,E) Representative flow cytometry histography showing Cy5 and GFP expression in HT1080 cells 48 h post-treatment with Cy5+EGFP-mRNA-NPs (1 nmol+10 µg/mL) or control-NPs. (C,D) Percentage of Cy5 and GFP in HT1080 cells (Control-Non treated) quantified by flow cytometry 48 h post-treatment with Cy5+EGFP-mRNA-NPs (1 nmol+10 µg/mL) or control-NPs (10 µM); n=3 (Data represent means ± SD, **** p < 0.0001). (F) Fluorescence microscopy images of HT1080 cells after 48 h treatment with dual-drug NPs or Empty NPs (scale bar-100 µm).

Next, we explored whether delivered mRNA and siRNA were functional and simultaneously processed by the cellular machinery to perform their intended functions. To achieve this, we developed monoclonal cell lines of MDA-MB-231 and HT1080 that stably express GFP, termed as MDA-MB-231-GFP^+^ and HT1080-GFP^+^ throughout the manuscript. These cell lines were created by retroviral transduction and subsequent ring cloning. Before we explored the transfection capacities of dual agent (M+S)-NPs, we studied single agent (M)-NPs and (S)-NPs. To this end, HT1080 and MDA-MB-231 cells were treated with (M)-NPs, and their corresponding GFP^+^ cell lines were treated with (S)-NPs for 48h and performed luminescence (Luciferase RLU) and flow cytometry assays, respectively. Our results showed that when compared to single agent (M)-NPs and (S)-NPs, they were successful in delivering functional agents in both cell lines, as shown in Figure 4 and Figure S7.

**Figure 4:**
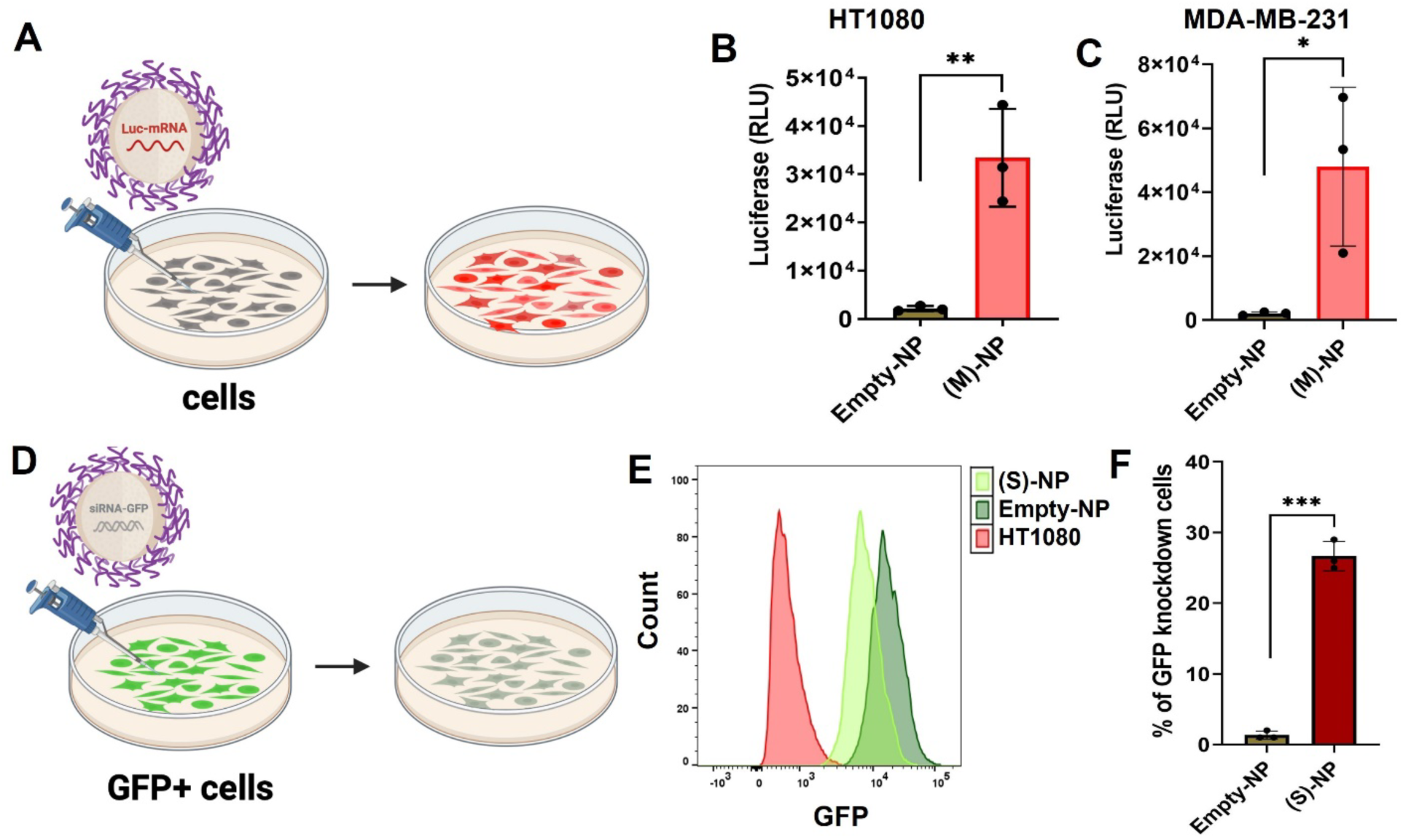
Single-drug NPs gene restoration and knock down. (A,B,C) Single drug (M)-NPs (10 µg/mL) and Empty-NPs (10 µM) treated to HT1080 and MDA-MB-231 cells and 48 h post-treatment luciferase (RLU) measured by plate reader n=3 (Data represent means ± SD, *p < 0.05 **p < 0.01). (D) Single drug (S)-NPs (10 µg/mL) and Empty-NPs (10 µM) treated to HT1080 GFP(+) cells. (E) Count of GFP knock down was analyzed by flow cytometry histogram; HT1080 cells used as positive control. (F) Percentage of GFP knock down was plotted with using flow cytometry analysis n=3 (Data represent means ± SD, ***p < 0.001).

To evaluate the transfection efficiency of our co-delivery NPs, we treated both GFP^+^ cells with (M+S)-NPs and control NPs for 48h. We assessed the simultaneous delivery of Luc-mRNA and siRNA-GFP by (M+S)-NPs and their effects on the cells. Flow cytometry was used to analyze GFP knockdown induced by siRNA-GFP from (M+S)-NPs, while a luciferase assay was performed to measure the expression of the luciferase gene by Luc-mRNA delivered *via* the same (M+S)-NPs. The results indicated that our (M+S)-NPs successfully escaped endosomes and simultaneously knocked down GFP expression with siRNA-GFP while introducing the expression of the luciferase gene via Luc-mRNA (**Figure 5B-C and Figure S7 & S8).** Compared to empty NPs, (M+S)-NPs produced a strong bioluminescence signal in both cell lines, confirming the successful delivery of Luc-mRNA by co-delivery NPs. Additionally, mRNA transfection efficiency was higher in HT1080-GFP+ than in MDA-MB-231-GFP+ cell lines. As expected, (M+S)-NPs-delivered siRNA-GFP also successfully knocked down GFP in both cell types, as evidenced by flow cytometry.

**Figure 5.**
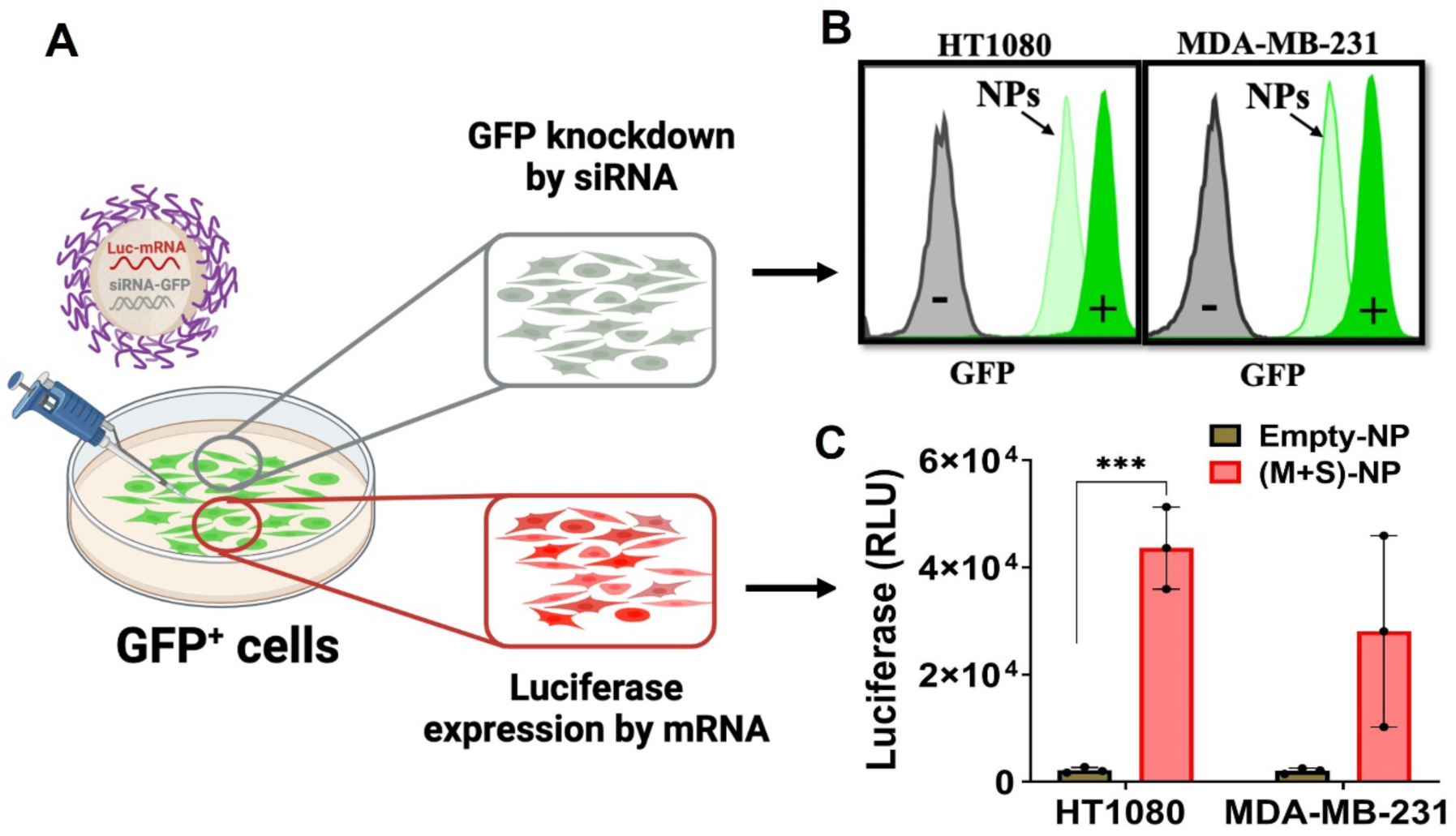
Simultaneous gene restoration and knockdown mediated by co-delivery NPs using mRNA and siRNA in two cell lines. (A) GFP+ cells (HT1080 and MDA-MB-231) were treated with (M+S)-NPs (∼60 nm size) co-loaded with siRNA-GFP (1pmol) and Luc mRNA (10 µg/mL) for 48h. (B) GFP knockdown count was analyzed via flow cytometry; GFP (-) cells used as positive control and (C) Luciferase expression was measured by luminescence n=3 (Data represent means ± SD, ***p < 0.001).

Overall, our *in vitro* studies with GFP^+^ cells showed that (M+S)-NPs can successfully escape the endosomes, simultaneously knocking down GFP expression while introducing luciferase gene expression into the cells.

### Codelivery NPs can express target genes within TNBC tumors while simultaneously silencing another gene *in vivo*

To assess the translation of our (M+S)-NPs’ *in vitro* effects into an *in vivo* setting, we studied their efficacy using a TNBC *in vivo* model. As a proof of concept, MDA-MB-231-GFP^+^ cells were implanted into the mammary gland of NSG mice to produce GFP^+^ TNBC tumors. Once tumors reached a mean diameter of 1cm, mice were randomized into two cohorts and treated with (M+S)-NPs and empty NPs for 24h.

Luciferase induction was examined using an In Vivo Imaging System (IVIS) at 0 and 20h post-treatment and with *ex vivo* bioluminescence at 24h. GFP knockdown was measured using flow cytometry after 24h of treatment. At 0h post-treatment, no luminescence was observed in either the control groups or (M+S)-NPs treated groups. However, at 20h post-treatment, a strong luciferase bioluminescence was detected in the tumors of mice treated with (M+S)-NPs, while those treated with empty NPs treatment showed no luminescence signal (Figure 6B). Subsequently, after the 24h treatment, tumors were extracted, single-cell suspensions were prepared, and luciferase bioluminescence and GFP were quantified using bioluminescence and flow cytometry, respectively. Compared to the control treatments, tumor cells treated with (M+S)-NPs exhibited a significantly strong (>300 folds) luciferase signal (Figure 6C). In addition, (M+S)-NPs achieved approximately a 20% knockdown of GFP expression in tumor cells (Figure 6D and E). These results unequivocally demonstrate that following administration, (M+S)-NPs can penetrate the tumor cells and deliver both mRNA and siRNA, enabling simultaneous knockdown of the target gene product while inducing expression of another protein.

**Figure 6.**
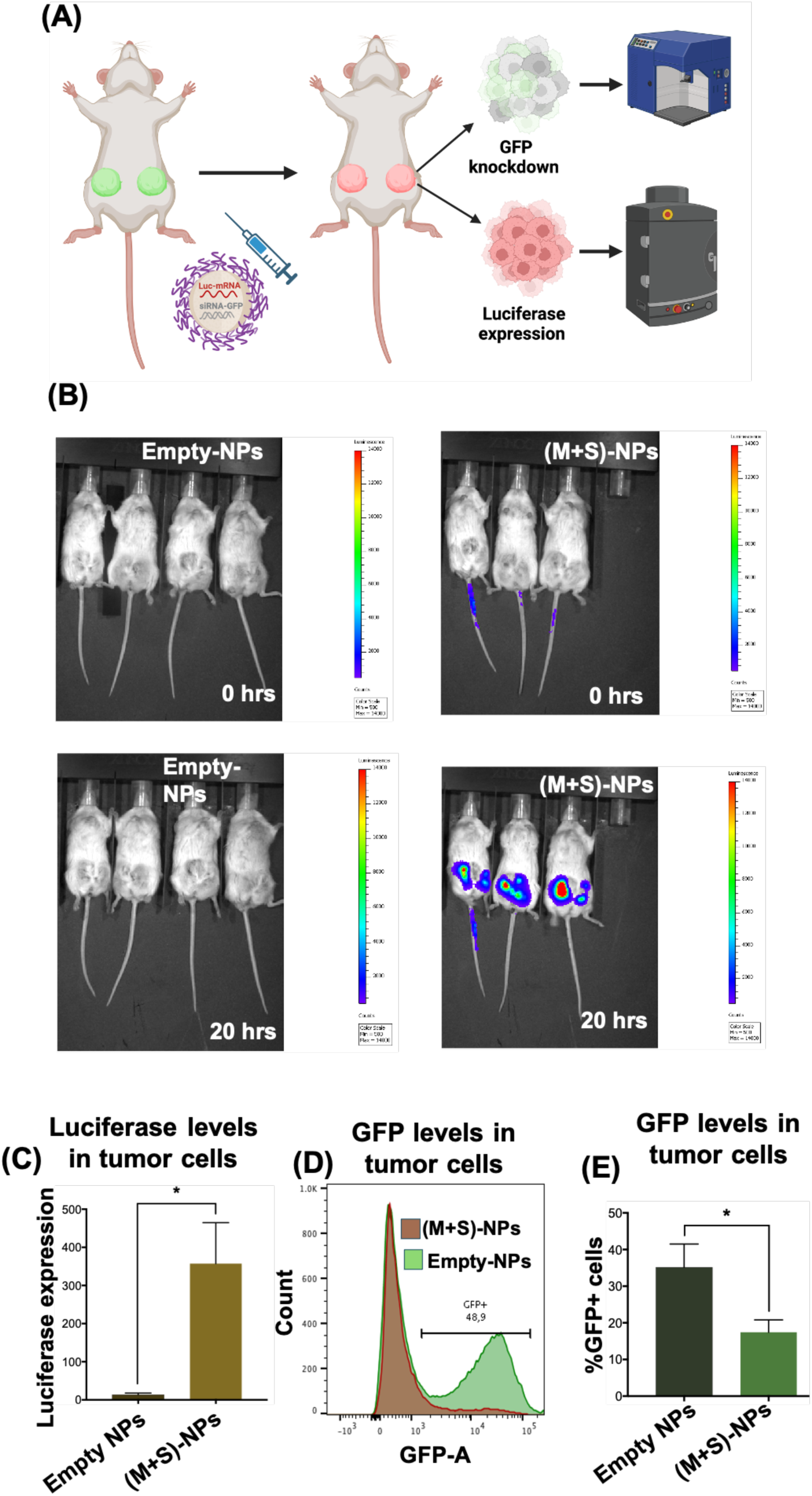
*In vivo*, co-delivery NPs achieve simultaneous functional restoration and gene knockdown using mRNA and siRNA. GFP+ cells (MDA-MB-231) implanted into the mammary gland of NSG mice were treated via intratumoral injection (IT) with (M+S)-NPs co-loaded with siRNA-GFP and Luc mRNA for 24 hours (A). Luciferase expression of Luc mRNA was measured using IVIS (B). GFP knockdown and luciferase expression were further analyzed via flow cytometry after dissociation of tumor into single cells (C and D), and percentage GFP+ cells were summarized (E). n=4 for Control-NPs and n=3 for (M+S)-NPs.

## Discussion

RNAi and mRNA technologies hold immense potential to treat a variety of diseases by targeting genes that are undruggable for traditional therapies.^4^ They offer versatile mechanisms for modulating gene/protein expression, including downregulation, augmentation, or correction. However, the full potential of RNA-based therapies remains unrealized due to inherent limitations, such as stability and unfavorable PK/PD properties. Effective delivery of RNA agents into cells is crucial for their treatment efficacy. In this context, advancements in NP-based drug delivery platforms have paved the way for novel NP systems capable of successfully delivering either siRNA or mRNA. Notable Current clinical success, such as Partisiran, a siRNA-based therapy for polyneuropathy in people with hereditary transthyretin-mediated amyloidosis, and mRNA-based COVID-19 vaccines, have accelerated RNA-based drug development for a variety of therapies.^8, 17^

In cancer, siRNA agents, alone or in combination with chemotherapeutic agents, are predominantly explored for their ability to silence proteins implicated in drug resistance.^13, 14^ Conversely, mRNA therapies are primarily studied for their potential to restore tumor suppressor gene/protein expressions, or to produce cytokines/antigens to improve anti-tumor immune response.^6^ However, cancer is a multifaceted disease, and genomic studies have highlighted that several factors are involved in cancer progression and survival.^14, 27^ Additionally, single-target therapies for cancer often encounter challenges such as drug resistance and tumorigenesis.^28^

In TNBC, conventional chemotherapeutic treatments are frequently associated with drug resistance, the development of cancer stem cells, tumorigenesis, and adverse side effects. Our previous research has shown that multi-targeted approaches can effectively control tumor growth and diminish CSC enrichment using clinically relevant tumor models.^21, 27^ In light of these findings, we reasoned that knocking down or restoring a single target gene may have limitations in effectively treating cancer, necessitating multi-targeted approaches. To address this, we developed (M+S)-NPs using siRNA-GFP and Luc-mRNA as model agents to simultaneously restore one gene while silencing the other. This approach represents a promising strategy for overcoming the challenges associated with single-target therapies and advancing the treatment of complex diseases like cancer.

Our results showed that (M+S)-NPs effectively encapsulated both agents, protected them from degradation, and successfully delivered them inside the cells. Notably, despite the size difference between siRNA (20-21 nucleotides) and Luc-mRNA (> 2000 nucleotides), the NPs efficiently encapsulated both within a single particle. Our NP’s sizes are around 60 nm comparable to our previous NPs known to accumulate in tumors after intravenous administration. ^27^ *In vitro* and *in vivo* characterization of (M+S)-NPs confirmed their ability to effectively enter cells, escape endosomes, and release their cargo in the cytosol. Additionally, (M+S)-NPs delivered siRNA-GFP and Luc-mRNA were functional and effective, as evidenced by a significant decrease in GFP expression and an increase in Luciferase signal in both *in vitro* and *in vivo* studies (Figure 5 and 6). Within the cells, the two RNA agents are processed differently. While mRNA is recruited into ribosomes for translation, siRNA is cooperated into the multiprotein RNA-induced silencing complex (RISC), which recognizes and cleaves complementary mRNA.^5, 12^ Our NPs results showed that these processes did not interfere with each other and operated independently. However, further studies are needed to understand the kinetics and any complementary interactions.

Diseases like cancer and cardiovascular diseases are complex and require multi-targeted approaches. Our proof-of-concept study demonstrated the feasibility of precisely modulating the expressions of multiple genes simultaneously. Drug resistance, CSCs enhancement, and tumorigenesis are primary challenges in current TNBC therapies. Our *in vivo* studies using the TNBC cell line model suggest that we could induce tumor suppressor gene expression such as PTEN, P53, or tumor antigens for immunotherapy while simultaneously knocking down the genes/proteins involved in drug resistance/CSC developments.^19, 29^ Although GFP and luciferase are not directly interconnected, several signaling pathway crosstalk, such as MAPK/ERK and PI3K-Akt, Wnt and NF-kB, JAK/STAT and RAS/MAPK have been reported in tumors. Our approach can selectively enhance and/or interfere with these cross-talks to improve therapeutic outcomes. Additionally, our study suggests that we can synergistically promote anti-tumoral factors while minimizing pro-tumoral factors via mRNA introduction, RNA interference, translational inhibition, and/or translational repression. Overall, the studies presented here will pave the way for new therapeutic strategies for complex diseases like cancer, with significant implications for drug development.

## Conclusions

RNA-based strategies, such as RNAi and mRNA technologies, hold significant promise in treating major diseases, including cancer. Current studies primarily focus on introducing tumor suppressors using mRNA or knocking down proteins that cause drug resistance with siRNA. However, cancer’s complexity necessitates multi-targeted approaches. To address this, we developed co-delivery NPs using Luc-mRNA and siRNA-GFP as model agents. Our NPs efficiently transfected and successfully functioned in two different cancer cell lines *in vitro*. When administrated to a TNBC mouse model created using MDA-MB-231 GFP+ cell line, the co-delivery NPs efficiently knocked down GFP while simultaneously introducing luciferase. This proof-of-concept has significant implications in developing RNA-based multi-targeted therapies for complex diseases like cancer.

## Materials and Methods

### Materials and reagents

mPEG-PLGA was procured from PolySciTech., Polyethylenimine (PEI) and 1,2-epoxytetradecane were purchased from Sigma Aldrich, EGFP-mRNA & Luc-mRNA purchased were from GenScript Biotech, GFP-siRNA was purchased from integrated DNA technologies, Cy5-siRNA universal negative control was purchased from Sigma Aldrich, Centrifuge filters obtained from Pall corporation, DMSO solvent obtained from fisher scientific, Copper grids were obtained from Ted Pella Inc, UranyLess EM stain was procured from Electron Microscopy Sciences, RiboGreen assay reagent and Luciferase reagent kit were purchased from Thermo fisher scientific, Dulbecco’s modified eagle’s medium purchased from Wisent bioproducts, and Trypsin for cell splitting was obtained from corning Inc.

### Nanoparticles synthesis and characterization

All NPs were synthesized using the nanoprecipitation method by mixing PLGA-PEG, PEI-C_14_, and mRNA/siRNAs in appropriate ratios. To mRNA (50 µg/mL) or siRNA (4 nmol) in 1 mL sterile water, slowly added PEI-C_14_ lipid (0.1 mM (0.2 mM for dual drug NPs)) with 200 µL of DMSO and gently spun the mixture. Subsequently, a solution of mPEG-PLGA (0.2 mM (0.4 mM for dual drug NPs)) in 200 µL DMF was added dropwise, and spinning continued for 2 h (same concentration of PEI-C_14_ and mPEG-PLGA used for Empty NP synthesis). Then, NPs solutions were concentrated for 2 rounds for 10 min. The resulting NPs were characterized for size and surface charge and stored at 4° C. The particle size distribution and morphological appearance of NPs were examined under transmission electron microscopy. Briefly, 10 µL of sample was spread onto a Cu grid (300 mesh) for 30 seconds and allowed to stain using 10 µL of UranyLess. The grid was dried before visualizing under JEM-1400Plus Transmission Electron Microscope operated at 120 kV. The images were acquired on a FIJI ImageJ software. Also, NPs were tested for stability by incubating them for 6 h in 5% or 10 % of fetal bovine serum (FBS) at 37 °C and then measured the size. The efficiency (EE%) of mRNA in the NPs measured using RiboGreen (Thermo Fisher Scientific) assay as previously described.^30^ Briefly, single or dual-drug NPs (15 µL) and 1X TE buffer (235 µL) were mixed as one solution. From this NPs solution, three different volumes were taken in replicate manner in 96-well black well plate. Next, 1% Triton X-100 buffer was added to each sample. RNA standard’s solutions were prepared in 1XTE buffer in the same plate. After 10-15 minutes, fluorescent RiboGreen reagent was added to NPs solutions and RNA standard solutions. This resulting solutions fluorescence intensity was read at excitation, 490 nm; emission, 520 nm, using the plate reader BioTek Synergy Neo2 Hybrid Multimode. To measure the amount of mRNA and determine the encapsulation efficiency, a standard curve was employed.

For siRNA encapsulation efficiency (EE%), Cy5-labeled siRNA fluorescent intensity was measured. Briefly, in black 96-well plate, Cy5-siRNA standard solutions (6 concentrations) were placed with increasing concentrations. Each Cy5-siRNA concentration’s volume made up to 100 µL with DMSO. In the same plate, Cy5-siRNA-based single or dual-drug NPs (50 µL) and DMSO (50 µL) were mixed in other wells in replicate manner. Fluorescence intensity was read at excitation, 650 nm; emission, 670 nm, using the plate reader BioTek Synergy Neo2 Hybrid Multimode. A standard curve was made to determine the amount of siRNA and the efficiency of encapsulation.

### *In vitro* studies

The MDA-MB-231 breast cancer cell line and HT1080 cell line were purchased from the American Type Culture Collection (ATCC, Manassas, VA, USA), both cells were maintained in DMEM media supplemented with 10% FBS and 1% penicillin/streptomycin.

To generate the GFP+ MDA-MB-231, GFP+ HT1080 monoclonal cell lines, retroviral vectors encoding GFP were produced by co-transfection of HEK293T cells (ATCC) with MLV LTR-GFP, gagpol, and a plasmid encoding VSV-G (all kind gifts of Dr. James Cunningham, Brigham and Women’s Hospital) using the JetPrime (Polyplus-transfection) reagent following the manufacturer’s protocol. Supernatants containing the retroviral pseudotypes were filtered (0.45µm) and used to transduce MDA-MB-231 and HT1080 cells in the presence of polybrene (3 µg/mL). 48 to 72 hours later, transduced cells were seeded at very low density in 10 cm-dishes. Cells were incubated at 37°C until the generation of colonies which were selected based on GFP expression as visualized using the ZOE imager (Bio-Rad) and picked using cloning cylinders.

### GFP knockdown

GFP +ve cells were seeded in a 12-well plate, at 50,000 cells per well. At 60% confluency, cells were treated with (S)-NP (1 nmol) or (M+S)-NP (1 nmol+10 µg/mL) and incubated for 48 hours. Then, cells were trypsinized and centrifuged. The cell pellet was resuspended with 1X FACS buffer. GFP knockdown was assessed quantitively by using the BD LSRFortessa flow cytometer and data analyzed with FlowJo software.

### (Cy5+EGFP-mRNA)-NP Co-delivery to HT1080 cells

In a 12-well plate, HT1080 cells were seeded with density of 50,000 cells per well. After one day, cells were treated with (Cy5+EGFP-mRNA)-NP (1 nmol+10 µg/mL) and incubated for 48 hours. GFP production of live cell imaging was performed by fluorescent microscopy using the ZOE Fluorescent Cell Imager (Bio-Rad, CA, USA). Followed by, cells were trypsinized and centrifuged. The cell pellet was resuspended in FACS buffer. Percentage of GFP production and Cy5 cellular entry were assessed quantitively by using the BD LSRFortessa flow cytometer and data analyzed with FlowJo software.

### Luciferase assay

In white 96-well plate, at density of 7×10^3^ cells per well (GFP+ or GFP-) were seeded as triplicate before one day to treatment. Single (10 µg/mL) or dual-drug NPs (1 nmol+10 µg/mL) treated to cells with fresh media and incubation continued for 48 h. Then, medium was removed and each well added with 50 µL 1X luciferase lysis buffer and lysed the cells performed by single freeze-thaw cycle. Followed by, 50 µL luciferin reagent added to each well and initiated the luminescence reading using BioTek Synergy Neo2 Hybrid Multimode Reader.

### *In vivo* studies

The animal studies were approved by the Animal Care and Use Committee at the University of Ottawa (protocol # BMIe-4035). The MDA-MB-231-GFP+ breast cancer cells were mixed 1:1 with Matrigel and injected under aseptic conditions into the mammary fat pads (2 × 10^6^ cells per fat pad) of NSG mice. When the tumor reached a mean diameter of approximately 1cm, tumor-bearing NSG mice were treated via intra-tumoral injection (IT) of 50µL (M+S)-NPs ((2.5 nmol siRNA-GFP + 0.25 mg Luc mRNA)/mouse) or empty NPs. After 20 hrs post-injection, relative bioluminescent intensity in tumors and different organs in mice was quantified using the Perkin Elmer IVIS Spectrum In Vivo Imaging System (IVIS). Following IVIS, mice were humanely euthanized 24h post-injection. The tumors were harvested and minced using a scalpel and incubated in DMEM media containing collagenase/hyaluronidase (STEMCELL Technologies, #07912) at 37 °C for 60 minutes. Afterwards the solution was passed through a 40 µM nylon mesh for the creation of a single cell solution. GFP knockdown was analyzed via flow cytometry and luciferase expression was assessed by bioluminescence. n=4 for control empty-NPs and n=3 for mRNA/siRNA NPs.

### Flow cytometry

Dissociated cancer cells were filtered through a 40µm strainer and suspended in PBS supplemented with 2% FBS and 2mM EDTA. 1µL of mouse IgG (1mg/mL) was added and incubated at 4° for 10 minutes. Afterwards, the cells were resuspended in 1x binding buffer (eBioscience, San Diego, CA, USA) and incubated with Annexin-V (eBioscience) for 15 minutes at room temperature. The cells were then washed twice and 7-aminoactinomycin D (7-AAD, eBioscience, San Diego, CA) was added to exclude dead cells. Flow cytometry was performed on a Cyan-ADP 9 or the BD LSRFortessa. Data was analyzed with FlowJo software (Ashland, OR, USA).

### Statistical analysis

Data are represented as means +/-Standard deviation (SD) or standard Error (SE). Statistical significance was determined using ANOVA or student t-test wherever appropriate and reported as (*) for p < 0.05, (**) for p < 0.01, (***) for p < 0.001, (****) for p < 0.0001, and n for no significant difference. Unless otherwise stated, experiments have a minimum of three biological repeats.

## Ethics approval

The animal studies were approved on 12/20/2023 by the Animal Care and Use Committee (ACC) at the University of Ottawa (protocol # BMIe-4035).

## Author contributions

SM and SE contributed equally to the manuscript. All authors were involved in data collection, analysis, and final editing.

## Acknowledgment

The authors would like to thank Ardeshir Ariana for help with flow cytometry and Vera Tang from the uOttawa Flow Cytometry and Virometry core for technical support.

## Supporting Information

The Supporting Information contains PEI-C_14_ synthesis, characterization. Cy5+EGFP-mRNA-NPs characterization and flow cytometry analysis of Cy5 and EGFP expression in HT1080 cells. Flow cytometry analysis of GFP knockdown in HT1080 GFP+ cells.

## Reference

(1) Huang, X.; Kong, N.; Zhang, X.; Cao, Y.; Langer, R.; Tao, W. The landscape of mRNA nanomedicine. Nat Med 2022, 28 (11), 2273–2287. DOI: 10.1038/s41591-022-02061-1 From NLM Medline.

(2) Mendes, B. B.; Conniot, J.; Avital, A.; Yao, D.; Jiang, X.; Zhou, X.; Sharf-Pauker, N.; Xiao, Y.; Adir, O.; Liang, H.;, et al. Nanodelivery of nucleic acids. Nat Rev Methods Primers 2022, 2. DOI: 10.1038/s43586-022-00104-y From NLM PubMed-not-MEDLINE.

(3) Kulkarni, J. A.; Witzigmann, D.; Thomson, S. B.; Chen, S.; Leavitt, B. R.; Cullis, P. R.; van der Meel, R. The current landscape of nucleic acid therapeutics. Nat Nanotechnol 2021, 16 (6), 630–643. DOI: 10.1038/s41565-021-00898-0 From NLM Medline.

(4) Hopkins, A. L.; Groom, C. R. The druggable genome. Nat Rev Drug Discov 2002, 1 (9), 727–730. DOI: 10.1038/nrd892 From NLM Medline.

(5) Qin, S.; Tang, X.; Chen, Y.; Chen, K.; Fan, N.; Xiao, W.; Zheng, Q.; Li, G.; Teng, Y.; Wu, M.;, et al. mRNA-based therapeutics: powerful and versatile tools to combat diseases. Signal Transduct Target Ther 2022, 7 (1), 166. DOI: 10.1038/s41392-022-01007-w From NLM Medline.

(6) Liu, C.; Shi, Q.; Huang, X.; Koo, S.; Kong, N.; Tao, W. mRNA-based cancer therapeutics. Nat Rev Cancer 2023, 23 (8), 526–543. DOI: 10.1038/s41568-023-00586-2 From NLM Medline.

(7) Hou, X.; Zaks, T.; Langer, R.; Dong, Y. Lipid nanoparticles for mRNA delivery. Nat Rev Mater 2021, 6 (12), 1078–1094. DOI: 10.1038/s41578-021-00358-0 From NLM PubMed-not-MEDLINE.

(8) Buschmann, M. D.; Carrasco, M. J.; Alishetty, S.; Paige, M.; Alameh, M. G.; Weissman, D. Nanomaterial Delivery Systems for mRNA Vaccines. Vaccines (Basel) 2021, 9 (1). DOI: 10.3390/vaccines9010065 From NLM PubMed-not-MEDLINE.

(9) Gao, M.; Tang, M.; Ho, W.; Teng, Y.; Chen, Q.; Bu, L.; Xu, X.; Zhang, X. Q. Modulating Plaque Inflammation via Targeted mRNA Nanoparticles for the Treatment of Atherosclerosis. ACS Nano 2023, 17 (18), 17721–17739. DOI: 10.1021/acsnano.3c00958 From NLM Medline.

(10) Parayath, N. N.; Stephan, S. B.; Koehne, A. L.; Nelson, P. S.; Stephan, M. T. In vitro-transcribed antigen receptor mRNA nanocarriers for transient expression in circulating T cells in vivo. Nat Commun 2020, 11 (1), 6080. DOI: 10.1038/s41467-020-19486-2 From NLM Medline.

(11) Kulkarni, J. A.; Witzigmann, D.; Chen, S.; Cullis, P. R.; van der Meel, R. Lipid Nanoparticle Technology for Clinical Translation of siRNA Therapeutics. Accounts of Chemical Research 2019, 52 (9), 2435–2444. DOI: 10.1021/acs.accounts.9b00368.

(12) Hu, B.; Zhong, L.; Weng, Y.; Peng, L.; Huang, Y.; Zhao, Y.; Liang, X. J. Therapeutic siRNA: state of the art. Signal Transduct Target Ther 2020, 5 (1), 101. DOI: 10.1038/s41392-020-0207-x From NLM Medline.

(13) Scully, M. A.; Wilhelm, R.; Wilkins, D. E.; Day, E. S. Membrane-Cloaked Nanoparticles for RNA Interference of beta-Catenin in Triple-Negative Breast Cancer. ACS Biomater Sci Eng 2024, 10 (3), 1355–1363. DOI: 10.1021/acsbiomaterials.4c00160 From NLM Medline.

(14) Gadde, S. Multi-drug delivery nanocarriers for combination therapy. MedChemComm 2015, 6 (11), 1916–1929, 10.1039/C5MD00365B. DOI: 10.1039/C5MD00365B.

(15) Paunovska, K.; Loughrey, D.; Dahlman, J. E. Drug delivery systems for RNA therapeutics. Nat Rev Genet 2022, 23 (5), 265–280. DOI: 10.1038/s41576-021-00439-4 From NLM Medline.

(16) Poon, W.; Kingston, B. R.; Ouyang, B.; Ngo, W.; Chan, W. C. W. A framework for designing delivery systems. Nat Nanotechnol 2020, 15 (10), 819–829. DOI: 10.1038/s41565-020-0759-5 From NLM Medline.

(17) Akinc, A.; Maier, M. A.; Manoharan, M.; Fitzgerald, K.; Jayaraman, M.; Barros, S.; Ansell, S.; Du, X.; Hope, M. J.; Madden, T. D.;, et al. The Onpattro story and the clinical translation of nanomedicines containing nucleic acid-based drugs. Nat Nanotechnol 2019, 14 (12), 1084–1087. DOI: 10.1038/s41565-019-0591-y From NLM Medline.

(18) Shi, J.; Kantoff, P. W.; Wooster, R.; Farokhzad, O. C. Cancer nanomedicine: progress, challenges and opportunities. Nat Rev Cancer 2017, 17 (1), 20–37. DOI: 10.1038/nrc.2016.108 From NLM Medline.

(19) Islam, M. A.; Xu, Y.; Tao, W.; Ubellacker, J. M.; Lim, M.; Aum, D.; Lee, G. Y.; Zhou, K.; Zope, H.; Yu, M.;, et al. Restoration of tumour-growth suppression in vivo via systemic nanoparticle-mediated delivery of PTEN mRNA. Nat Biomed Eng 2018, 2 (11), 850–864. DOI: 10.1038/s41551-018-0284-0 From NLM Medline.

(20) Xu, X.; Saw, P. E.; Tao, W.; Li, Y.; Ji, X.; Yu, M.; Mahmoudi, M.; Rasmussen, J.; Ayyash, D.; Zhou, Y.;, et al. Tumor Microenvironment-Responsive Multistaged Nanoplatform for Systemic RNAi and Cancer Therapy. Nano Lett 2017, 17 (7), 4427–4435. DOI: 10.1021/acs.nanolett.7b01571 From NLM Medline.

(21) El-Sahli, S.; Hua, K.; Sulaiman, A.; Chambers, J.; Li, L.; Farah, E.; McGarry, S.; Liu, D.; Zheng, P.; Lee, S. H.;, et al. A triple-drug nanotherapy to target breast cancer cells, cancer stem cells, and tumor vasculature. Cell Death Dis 2021, 12 (1), 8. DOI: 10.1038/s41419-020-03308-w From NLM Medline.

(22) Kamaly, N.; Yameen, B.; Wu, J.; Farokhzad, O. C. Degradable Controlled-Release Polymers and Polymeric Nanoparticles: Mechanisms of Controlling Drug Release. Chem Rev 2016, 116 (4), 2602–2663. DOI: 10.1021/acs.chemrev.5b00346 From NLM Medline.

(23) Xu, X.; Xie, K.; Zhang, X. Q.; Pridgen, E. M.; Park, G. Y.; Cui, D. S.; Shi, J.; Wu, J.; Kantoff, P. W.; Lippard, S. J.;, et al. Enhancing tumor cell response to chemotherapy through nanoparticle-mediated codelivery of siRNA and cisplatin prodrug. Proc Natl Acad Sci U S A 2013, 110 (46), 18638–18643. DOI: 10.1073/pnas.1303958110 From NLM Medline.

(24) Riley, R. S.; Kashyap, M. V.; Billingsley, M. M.; White, B.; Alameh, M. G.; Bose, S. K.; Zoltick, P. W.; Li, H.; Zhang, R.; Cheng, A. Y.;, et al. Ionizable lipid nanoparticles for in utero mRNA delivery. Sci Adv 2021, 7 (3). DOI: 10.1126/sciadv.aba1028 From NLM Medline.

(25) Smith, T. K. T.; Kahiel, Z.; LeBlond, N. D.; Ghorbani, P.; Farah, E.; Al-Awosi, R.; Cote, M.; Gadde, S.; Fullerton, M. D. Characterization of Redox-Responsive LXR-Activating Nanoparticle Formulations in Primary Mouse Macrophages. Molecules 2019, 24 (20). DOI: 10.3390/molecules24203751 From NLM Medline.

(26) Zhu, X.; Xu, Y.; Solis, L. M.; Tao, W.; Wang, L.; Behrens, C.; Xu, X.; Zhao, L.; Liu, D.; Wu, J.;, et al. Long-circulating siRNA nanoparticles for validating Prohibitin1-targeted non-small cell lung cancer treatment. Proc Natl Acad Sci U S A 2015, 112 (25), 7779–7784. DOI: 10.1073/pnas.1505629112 From NLM Medline.

(27) Sulaiman, A.; McGarry, S.; El-Sahli, S.; Li, L.; Chambers, J.; Phan, A.; Al-Kadi, E.; Kahiel, Z.; Farah, E.; Ji, G.;, et al. Nanoparticles Loaded with Wnt and YAP/Mevalonate Inhibitors in Combination with Paclitaxel Stop the Growth of TNBC Patient-Derived Xenografts and Diminish Tumorigenesis. Advanced Therapeutics 2020, 3 (11), 2000123. DOI: 10.1002/adtp.202000123.

(28) Sulaiman, A.; McGarry, S.; El-Sahli, S.; Li, L.; Chambers, J.; Phan, A.; Cote, M.; Cron, G. O.; Alain, T.; Le, Y.;, et al. Co-targeting Bulk Tumor and CSCs in Clinically Translatable TNBC Patient-Derived Xenografts via Combination Nanotherapy. Mol Cancer Ther 2019, 18 (10), 1755–1764. DOI: 10.1158/1535-7163.MCT-18-0873 From NLM Medline.

(29) Kong, N.; Tao, W.; Ling, X.; Wang, J.; Xiao, Y.; Shi, S.; Ji, X.; Shajii, A.; Gan, S. T.; Kim, N. Y.;, et al. Synthetic mRNA nanoparticle-mediated restoration of p53 tumor suppressor sensitizes p53-deficient cancers to mTOR inhibition. Sci Transl Med 2019, 11 (523). DOI: 10.1126/scitranslmed.aaw1565 From NLM Medline.

(30) Billingsley, M. M.; Singh, N.; Ravikumar, P.; Zhang, R.; June, C. H.; Mitchell, M. J. Ionizable Lipid Nanoparticle-Mediated mRNA Delivery for Human CAR T Cell Engineering. Nano Lett 2020, 20 (3), 1578–1589. DOI: 10.1021/acs.nanolett.9b04246 From NLM Medline.

