## Supplementary material for "Nanoparticles co-delivering siRNA and mRNA for simultaneous restoration and silencing of gene/protein expression *in vitro* and *in vivo*": SI

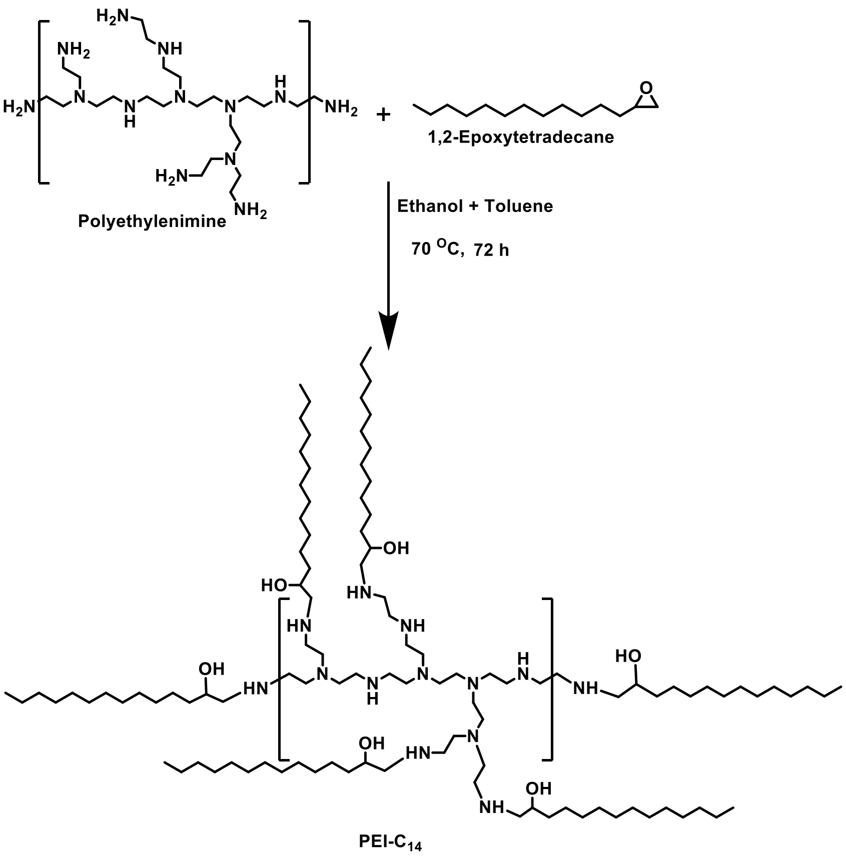


**Figure S1.** PEI-$C_{14}$ lipid synthetic route: epoxide ring opening by polyethylenimine in presence of ethanol and toluene solvents at 70 °C.

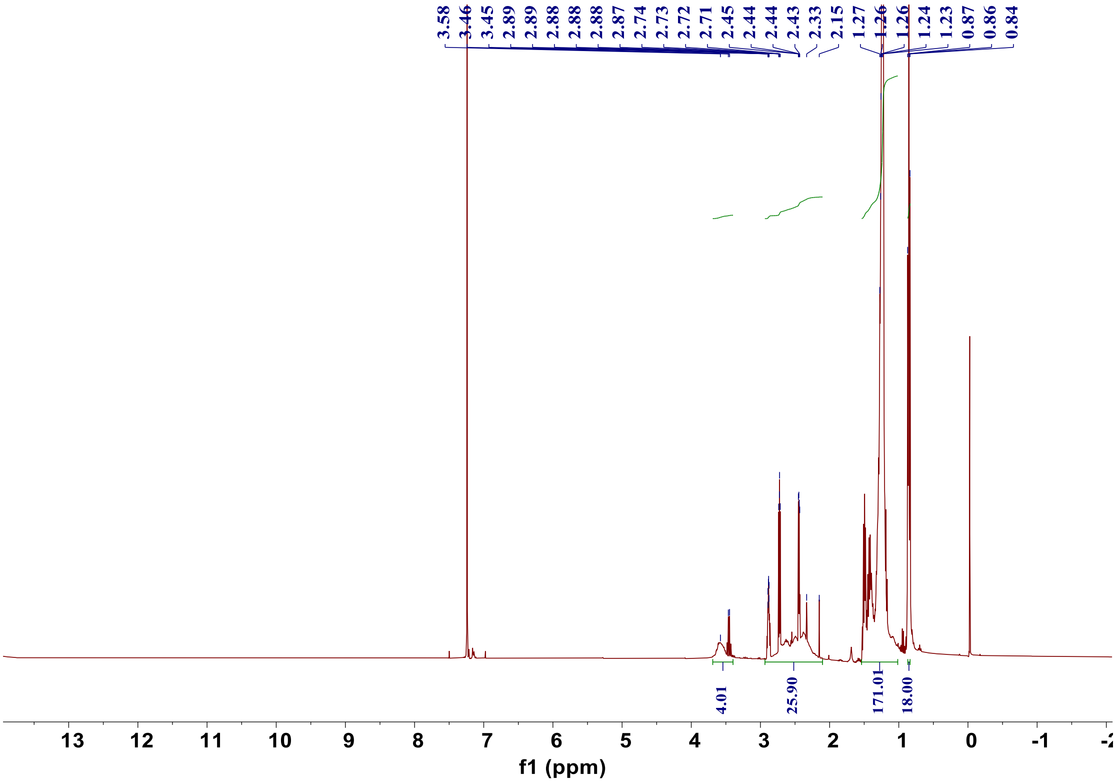


**Figure S2.** $^{1}$H (400 MHz Bruker AVANCE) NMR Spectrum of PEI-$C_{14}$ lipid; $^{1}$H NMR (400 MHz, $CDCl_3$) δ 3.71 – 3.30 (m, 4H), 2.91 – 2.10 (m, 26H), 1.51-1.24 (d, m 171H), 0.92 – 0.76 (m, 18H).

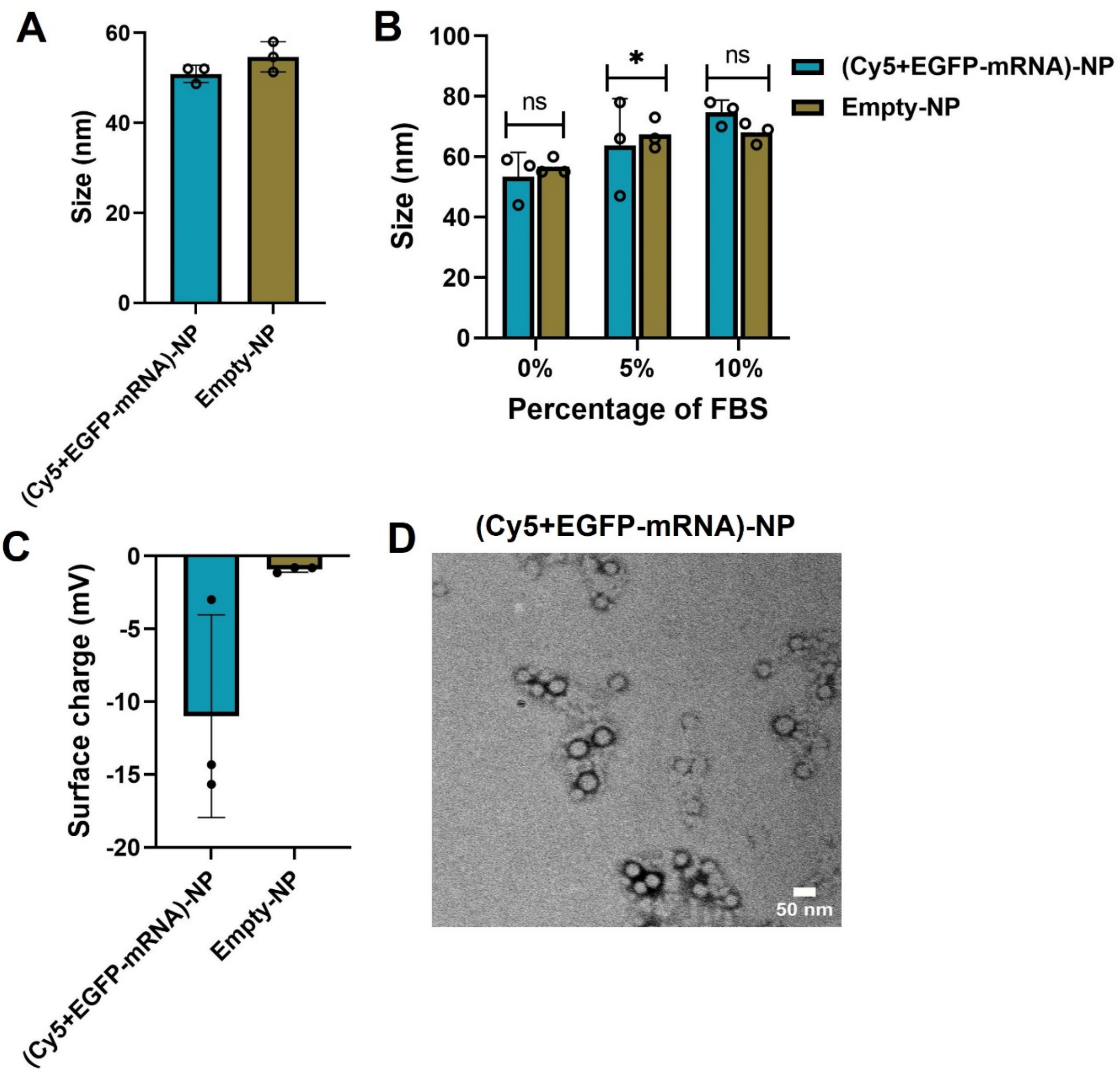


**Figure S3.** (A) Size of Cy5+EGFP-mRNA-NPs along with control-NPs, measured by diluting 20 µL of NP in 1 mL sterile water using Dynamic light scattering n=3. (B) Stability of Cy5+EGFP-mRNA-NPs in different percentage of FBS; NPs incubated for 6 hours and measured the size; n=3 (ns-no significant difference). (C) Surface of Cy5+EGFP-mRNA-NPs along with control-NPs, measured by diluting 20 µL of NP in 1 mL using Dynamic light scattering n=3. (D) Size and morphology of Cy5+EGFP-mRNA-NPs by transmission electron microscopy (scale bar-50 nm).

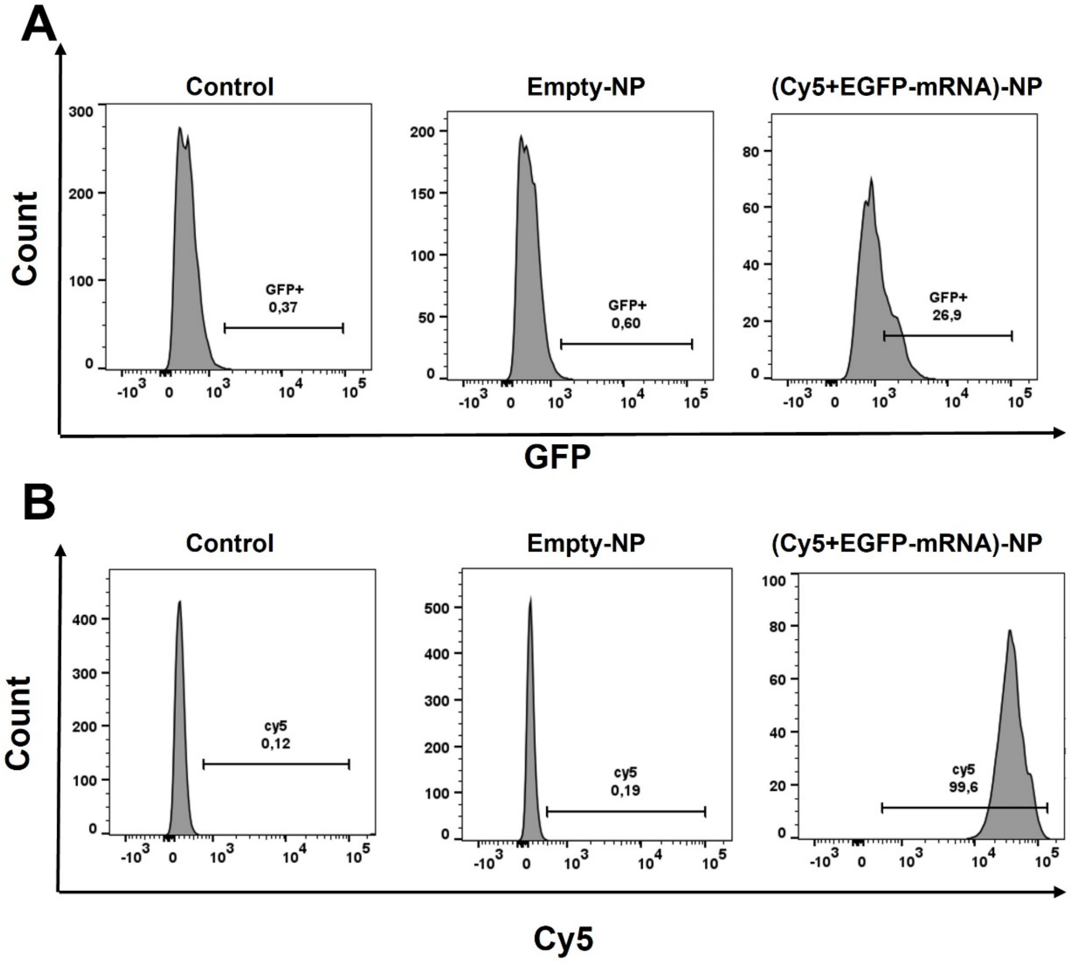


**Figure S4.** Representative flow cytometry histogram showing GFP (A) and Cy5 (B) expression in HT1080 cells after treated with (48 h post-treatment) Cy5+EGFP-mRNA-NPs and Empty-NPs (control is non-treated).

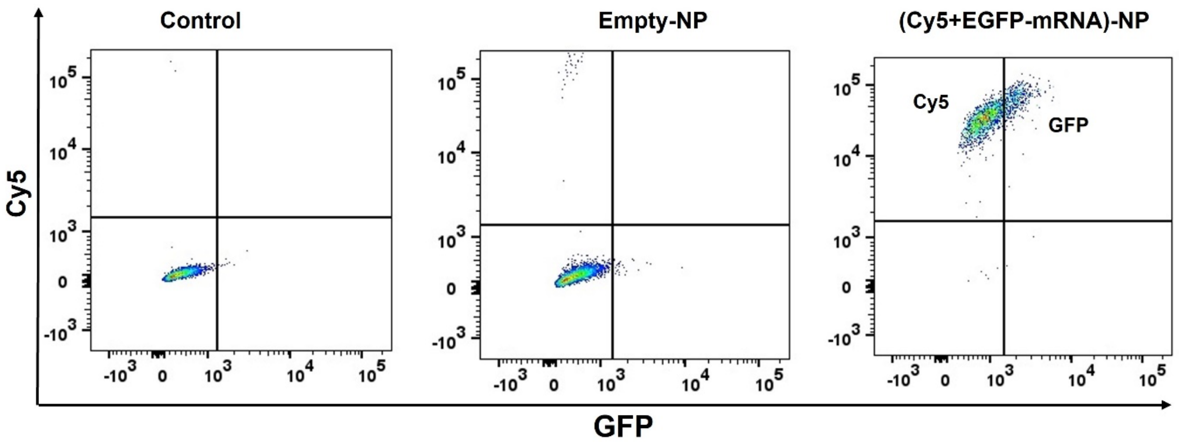


**Figure S5.** Cy5+EGFP-mRNA-NPs or Empty-NPs treated to HT1080 cells. Representative flow cytometry histogram showing dual-drug in single plot, GFP population in x-axis and Cy5 population in y-axis; control is non-treated.

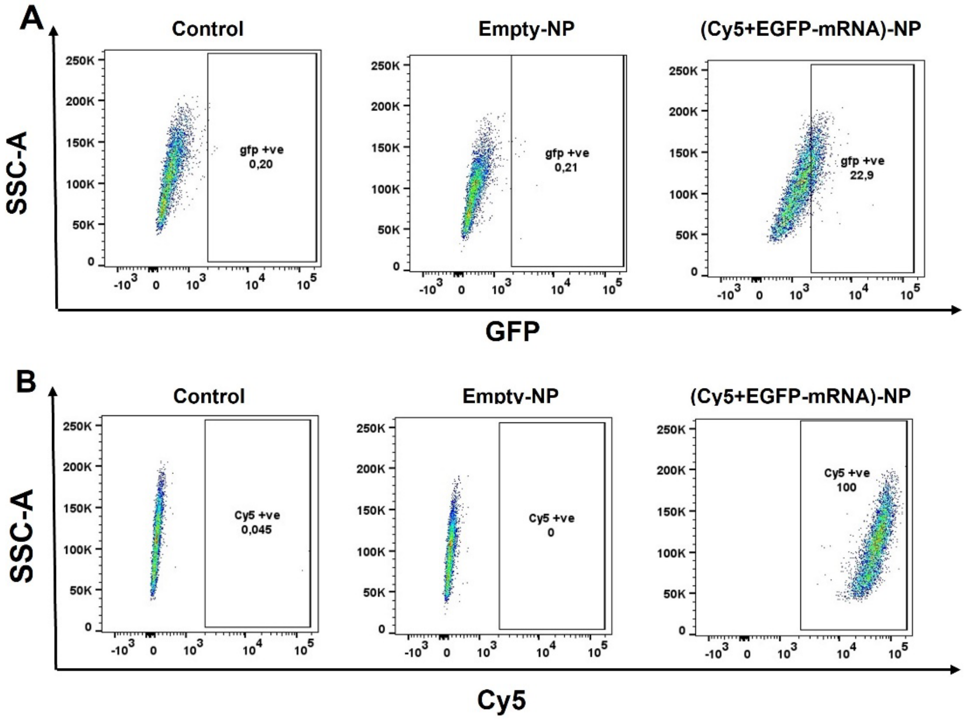


**Figure S6.** Representative flow cytometry SSC-A plots showing GFP (A), and Cy5 (B) expression in HT1080 cells 48 h post-treatment with Cy5+EGFP-mRNA-NPs and Empty-NPs (control are non-treated).

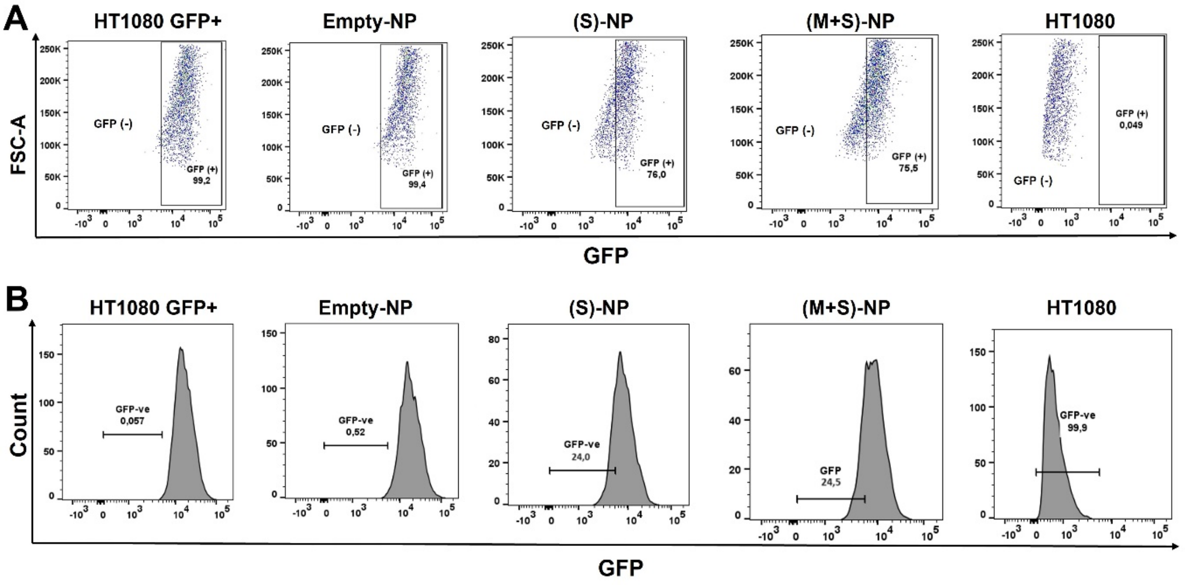


**Figure S7.** Single or dual-NPs along with Empty-NPs treated to HT1080 GFP (+) cells and 48 h post-treatment, count of GFP knock down was quantified by flow cytometry; (A) FSC-A plots; (B) histograms representing count of GFP knock down cells**.** HT1080 cells used as positive control.

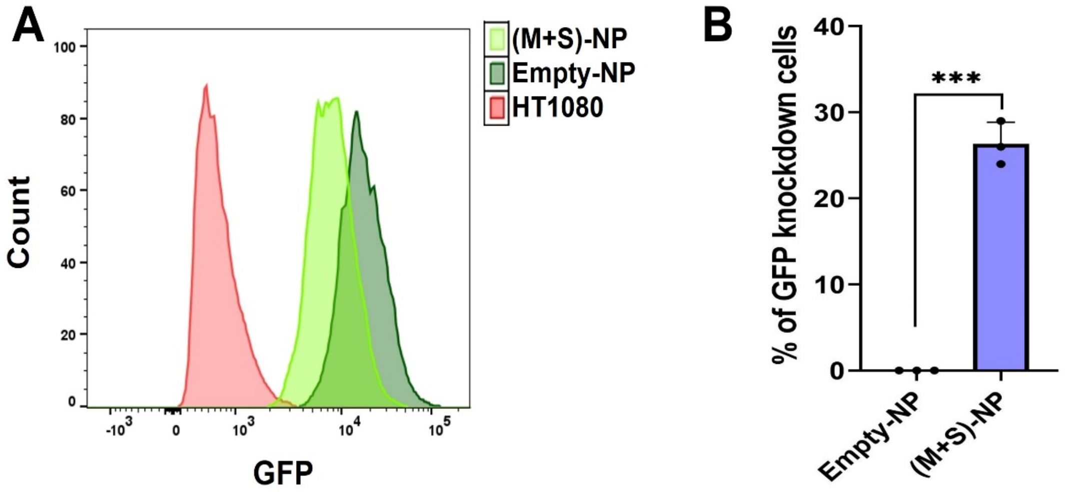


**Figure S8.** (A) Flow cytometry histogram for GFP knockdown with dual-drug-NPs in HT080 GFP(+) cells; HT180 cells used as positive control. (B) Percentage of GFP knock down was quantified with flow cytometry analysis; n=3 (Data represent means ± SD, ***$p < 0.001$).
